# Pharmacological elevation of cellular dihydrosphingomyelin provides a novel antiviral strategy against West Nile virus infection

**DOI:** 10.1101/2022.12.19.521146

**Authors:** Nereida Jiménez de Oya, Ana San-Félix, Mireia Casasampere, Ana-Belén Blázquez, Patricia Mingo-Casas, Estela Escribano-Romero, Eva Calvo-Pinilla, Teresa Poderoso, Josefina Casas, Juan-Carlos Saiz, María-Jesús Pérez-Pérez, Miguel A. Martín-Acebes

## Abstract

Flavivirus life cycle is strictly dependent on cellular lipid metabolism. Polyphenols like gallic acid and its derivatives are promising lead compounds for new therapeutic agents as they can exert multiple pharmacological activities, including the alteration of lipid metabolism. The evaluation of our own collection of polyphenols against West Nile virus, a representative medically relevant flavivirus, led to the identification of *N,N*′-(dodecane-1,12-diyl)bis(3,4,5-trihydroxybenzamide) and its 2,3,4-trihydroxybenzamide regioisomer as selective antivirals with low cytotoxicity and high antiviral activity (EC_50_ of 2.2 and 0.24 μM, respectively in Vero cells; EC_50_ of 2.2 and 1.9 μM, respectively in SH-SY5Y cells). These polyphenols also inhibited the multiplication of other flaviviruses, namely Usutu, dengue, and Zika viruses, exhibiting lower antiviral or negligible antiviral activity against other RNA viruses. The mechanism underlying their antiviral activity against WNV involved the alteration of sphingolipid metabolism. These compounds inhibited ceramide desaturase (Des1) promoting the accumulation of dihydrosphingomyelin (dhSM), a minor component of cellular sphingolipids with important roles on membrane properties. Addition of exogenous dhSM, or Des1 blockage by using the reference inhibitor GT-11, confirmed the involvement of this pathway in WNV infection. These results unveil the potential of novel antiviral strategies based on the modulation of the cellular levels of dhSM and Des1 activity for the control of flavivirus infection.

## Introduction

Polyphenols are secondary metabolites widely distributed in all higher plants which have important defensive roles against pathogenic microbes and herbivores, and serve to protect plants from several environmental stresses, such as rainfall and ultraviolet radiation (1). They are also responsible for flower coloration (2). Over 8000 polyphenols have been identified until now (3). They are organized in different families according to their structures, being phenolic acids one of the most representatives (4). To this group belongs gallic acid (3,4,5-trihydroxybenzoic acid), a low molecular weight triphenol abundantly present in free or ester forms in plants, and also in different vegetable foods such as grapes, nuts, berries, etc. and in beverages as wine, coffee and tea (5). Gallic acid and its esters (gallates) have shown preventive and therapeutic effects in many diseases where the oxidative stress has been involved, including cancer (6), cardiovascular diseases (7), neurodegenerative disorders (8) and aging (9). In addition, many antimicrobial activities such as antibacterial (10), antifungal (11) and antiviral (12–16) have been described for this family of polyphenols. Due to this, gallic acid and its derivatives are considered as promising lead compounds for new therapeutic agents (17). Our research group has synthesized alkyl esters and amides of gallic acid and its regioisomer, 2,3,4-trihydroxybenzoic acid. These compounds displayed antiviral activity against human hepatitis C virus (HCV), a single–stranded RNA virus belonging to the genus *Hepacivirus* within the family *Flaviviridae*. Preliminary data suggested that the antiviral effect against HCV of our galloyl derivatives could be related to alterations of cellular lipids (18) although the specific mechanism was not elucidated.

The family *Flaviviridae* also includes arthropod-borne pathogens transmitted by mosquitos or ticks classified in the genus *Flavivirus*, which groups more than 50 species (19). Vector-borne flaviviruses are responsible for a variety of human and animal illnesses (20). West Nile virus (WNV) and Usutu virus (USUV) cause neurological diseases (21, 22), yellow fever virus and dengue virus (DENV) cause hemorrhagic fevers (23), and Zika virus (ZIKV) is responsible of birth defects and an autoimmune disease (Guillain-Barré syndrome) with neurological symptoms (24). In a similar manner to that observed for other arthropod-borne viruses (arboviruses), the impact of flaviviruses in human and animal health has increased during the last decades due to a variety of factors that include climate warming, globalization of travel and trade, changes in land use and urbanization and alterations of vector behaviors (20, 25). Nowadays, there is no specific therapy against any flavivirus and only a few preventive vaccines are available. Therefore, the need for new antiviral therapies is a priority to combat these pathogens. Among antiviral strategies under investigation, host-targeted antivirals have raised as feasible alternatives to combat viral infections in contrast to specific antivirals targeting viral components (26). Their theoretical advantages include their potential broad-spectrum against present and future viral threats that share dependency on host factors and a high genetic barrier for the development of viral resistance (27, 28). In this scenario, growing evidences support the druggability of lipid metabolism to inhibit viral infections (29, 30). In the case of flaviviruses, infection is strictly dependent on specific lipids required for intracellular membrane rearrangements to establish viral replication factories, energy metabolism, innate immune evasion and particle biogenesis (31). In fact, certain sphingolipids, such as ceramide (Cer) and sphingomyelin (SM), play crucial roles in flavivirus infection (32–34).

We have evaluated the antiviral potential of a series of polyphenols in the infection of WNV, here selected as a representative medically relevant flavivirus. Our results indicate that these compounds reduce WNV multiplication by altering sphingolipid metabolism due to inhibition of Cer desaturase (Des1), leading to the accumulation of dihydrosphingomyelin (dhSM), a minor component of cellular sphingolipids with important roles on membrane properties. Consistent with the broad-spectrum antiviral potential of polyphenols (35), these compounds also inhibited the multiplication of the clinically relevant related flaviviruses USUV, DENV and ZIKV. Overall, these results highlight the potential of synthetic polyphenols for the development of broad-spectrum antiviral therapies and unveil a novel antiviral strategy against flaviviruses by intervention on the cellular levels of dhSM.

## Results

### Antiviral activity of gallic acid derivatives against WNV

The structure of the compounds under study are shown in Fig. 1A and include gallic acid, the octyl ester of gallic acid (AL-071), esters AL-072 and AL-085, containing an octyl or dodecyl methylene linker connecting two subunits of gallic acid at the ends, and amides AL-088 and AL-274, containing a dodecyl methylene linker and two galloyl moieties (AL-088) or its 2,3,4-trihydroxybenzoyl isomeric ring respectively at both ends (AL-274). The antiviral activity of these compounds against WNV in Vero cells, together with their effect on cell viability, estimated by ATP measurement, is shown in Fig. 1B. Values of the virus yield in PFU/mL are shown in supplementary Table 1. Table 1 summarizes the half-maximal effective concentrations (EC_50_), half-maximal cytotoxic concentration (CC_50_) and selectivity indexes (SI) of these compounds. Gallic acid showed a slight reduction in WNV yield concomitant with the reduction on cellular viability suggesting that the effect on virus growth was due to cytotoxicity rather than to a specific antiviral activity. The introduction of an acyl chain containing 8 methylenes resulted in the alkyl gallate AL-071 that showed some antiviral activity, but also higher cytotoxicity. The incorporation of a second gallic unit at the distal position of the octyl spacer (compound AL-072) did not improve the antiviral efficacy. However, when the linker between both gallic acid units was extended to a dodecyl unit (AL-085) the antiviral efficacy was significantly improved. Even more, the substitution of ester bonds to amides between the galloyl or 2,3,4-trihydroxybenzoyl units and the dodecyl methylene linker gave compounds AL-088 and AL-274 that resulted in significant antiviral activity against WNV (EC_50_ values in the low μM range) and greatly reduced cytotoxicity conferring remarkable selectivity indexes. In addition, the presence of amide bonds linking the distal polyphenols with the polymethylene chain should also improve the metabolic stability versus esterases. AL-088 and AL-274 exerted a limited impact on the proliferation capability of Vero cells, further supporting their low cytotoxic profiles (Fig. 1C). The antiviral activity of AL-088 and AL-274 was very similar despite of the MOI used in the experiments (Fig. 1D). As the central nervous system is a major target for WNV replication, the antiviral activity of compounds AL-088 and AL-274 was evaluated in SH-SY5Y cells, a human neuroblastoma cell line susceptible to WNV infection (46). Both compounds exhibited potent antiviral activity (EC_50_ values of 2.2 and 1.9 for AL-088 and AL-274, respectively), retaining good selectivity indexes (Fig. 2 and Table 1). Therefore, compounds AL-088 and AL-274 were advanced for further studies to evaluate their antiviral properties.

**Figure 1.**
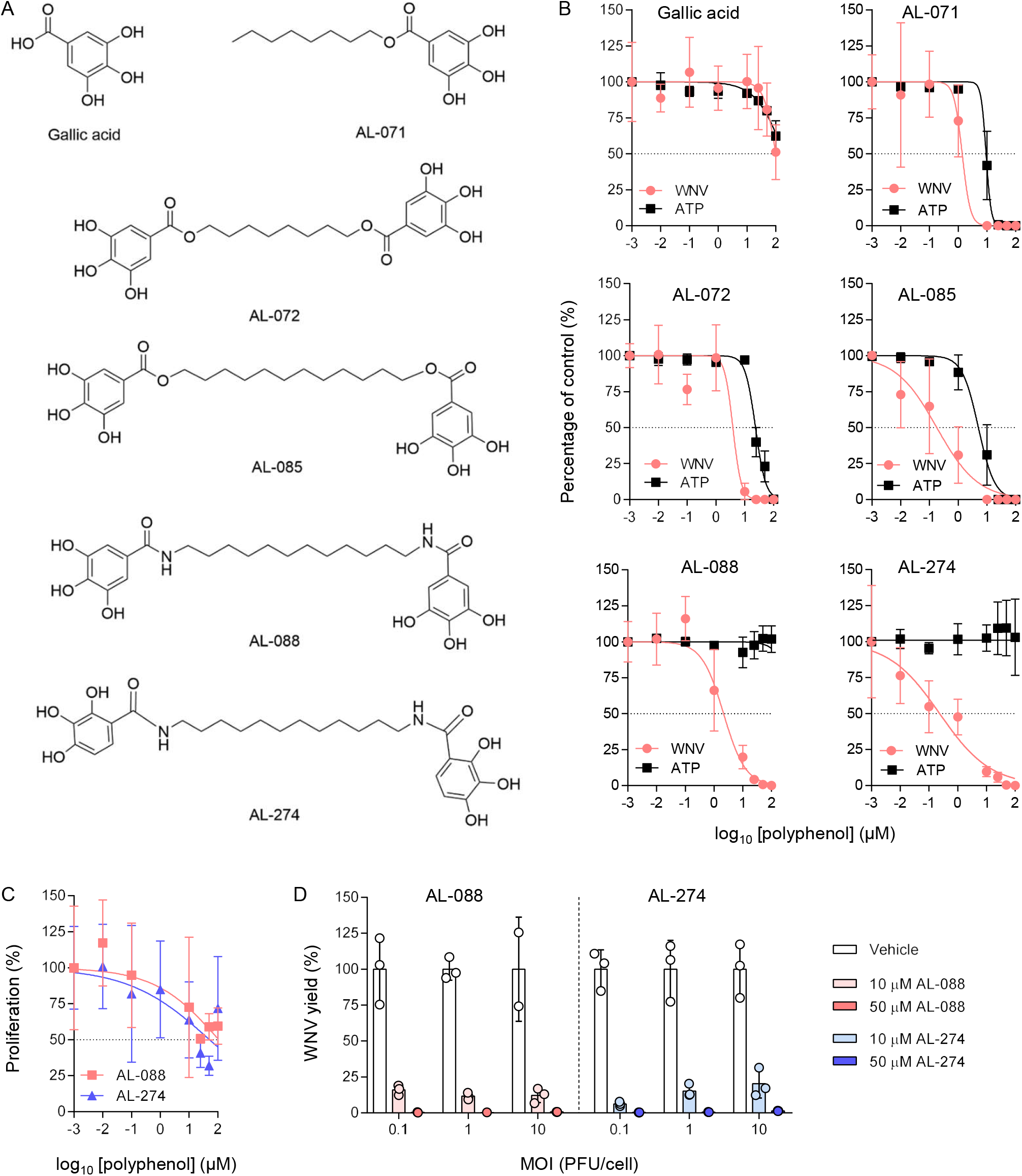
Antiviral activity of synthetic polyphenols against WNV in Vero cells. (A) Compounds used in this study. (B) Antiviral activity against WNV and cytotoxicity of polyphenols. Virus yield in the supernatant of infected Vero cells (MOI of 1 PFU/cell) was determined at 24 h pi. Polyphenols were added 1 h prior to infection and maintained throughout the rest of the assay. The cytotoxicity of the compounds was measured by determination of cellular ATP in uninfected samples. Dashed lines denote a 50% reduction. (C) Effect of polyphenols on the proliferation of Vero cells. Vero cells plated at low density (confluence <50%) were incubated for 24 h in the presence of the compounds and cell number was determined. Dashed lines denote a 50% reduction. (D) Effect of MOI on the antiviral activity of polyphenols. Vero cells were treated and infected at different MOIs as described in (A). Each dot denotes a single biological replicate. Data are expressed as mean ± SD (n = 2 to 4).

**Figure 2.**
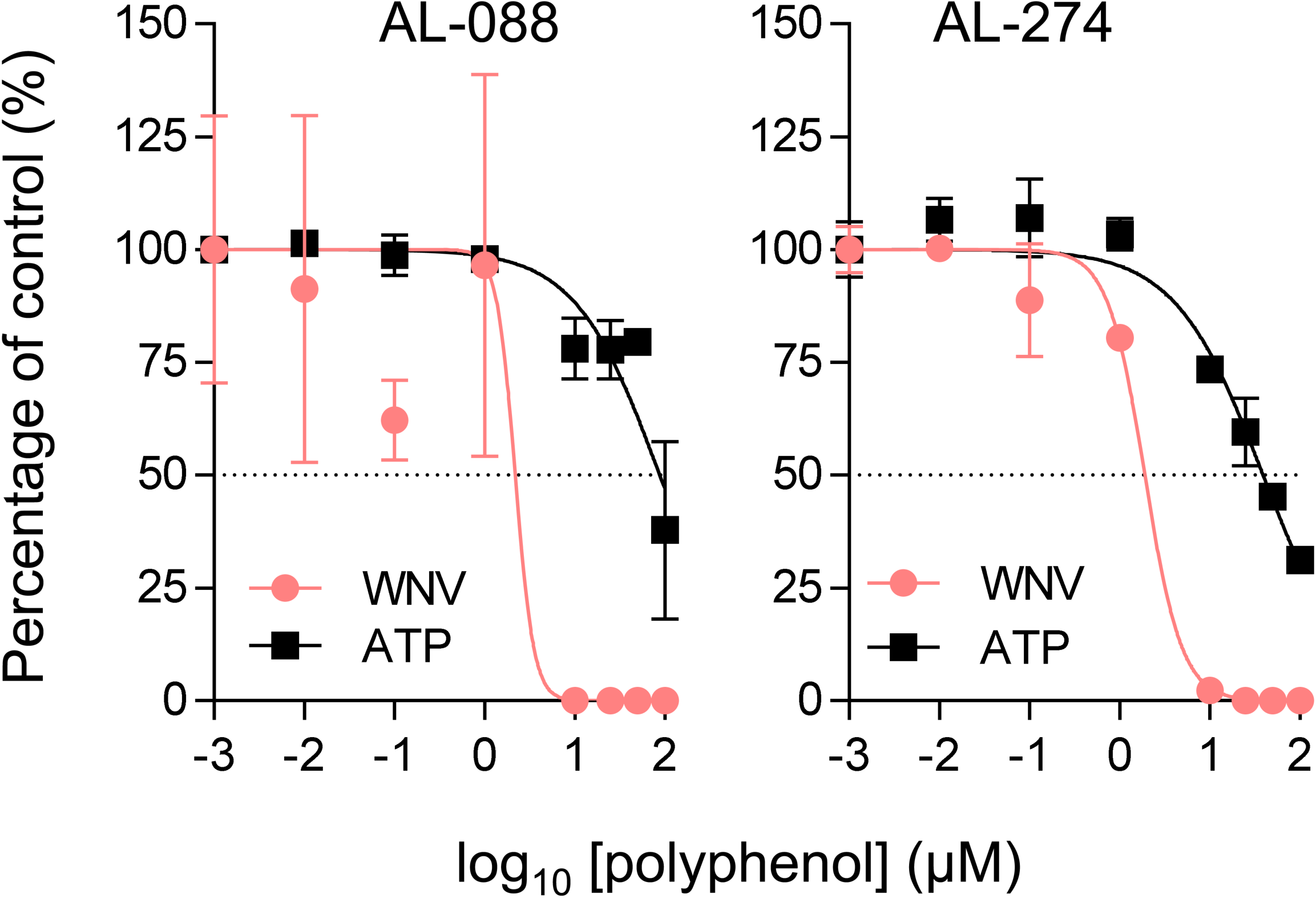
Antiviral activity of synthetic polyphenols against WNV in SH-SY5Y cells. Virus yield in the supernatant of infected SH-SY5Y cells (MOI of 1 PFU/cell) was determined at 24 h pi. Polyphenols were added 1 h prior to infection and maintained throughout the rest of the assay. The cytotoxicity of the compounds was measured by determination of cellular ATP in uninfected samples. Dashed lines denote a 50% reduction. Data are expressed as mean ± SD (n = 2 to 4).

**Table 1.** Antiviral efficacy and cytotoxicity of polyphenols.

|  | Virus | Vero |  |  |  |  |  | SH-SY5Y |  |
| --- | --- | --- | --- | --- | --- | --- | --- | --- | --- |
|  |  | Gallic acid | AL-071 | AL-072 | AL-085 | AL-088 | AL-274 | AL-088 | AL-0274 |
| CC<sub>50</sub><sup>a</sup><br>(μM) | Uninfected (24 h treatment) | >100 | 9.2 | 23.2 | 5.0 | >100 | >100 | 87.6 | 38.9 |
|  | Uninfected (48 h treatment) |  |  |  |  | >100 | >100 |  |  |
| EC<sub>50</sub><br>(μM) | WNV | ~100 | 1.4 | 3.9 | 0.2 | 2.2 | 0.24 | 2.2 | 1.9 |
|  | USUV |  |  |  |  | 0.8 | 1.3 |  |  |
|  | ZIKV |  |  |  |  | 6.7 | 2.1 |  |  |
|  | DENV-2 |  |  |  |  | 7.2 | 9.2 |  |  |
|  | VSV |  |  |  |  | 25.3 | 43.0 |  |  |
|  | CVB5 |  |  |  |  | >100 | >100 |  |  |
| SI<br>(CC<sub>50</sub>/<br>EC<sub>50</sub>) | WNV | >1.0 | 6.6 | 5.9 | 25.0 | >45.0 | >416 | 39.8 | 20.5 |
|  | USUV |  |  |  |  | >125 | >76.9 |  |  |
|  | ZIKV |  |  |  |  | >14.9 | >47.6 |  |  |
|  | DENV-2 |  |  |  |  | >13.9 | >10.8 |  |  |
|  | VSV |  |  |  |  | >4.0 | >2.3 |  |  |
|  | CVB5 |  |  |  |  | 1 | 1 |  |  |

<sup>a</sup> Data are expressed in μM and correspond to the mean (n=2-4)

### Antiviral spectrum of AL-088 and AL-274

Compounds AL-088 and AL-274 also showed potent antiviral activity against the flaviviruses USUV, ZIKV and DENV-2 (Fig. 3 and Table 1), especially remarkable against USUV with EC_50_ values of 0.8 and 1.3 μM, respectively. However, their antiviral efficacy was reduced against VSV, an RNA virus from the family *Rhabdoviridae*, or even disappear against CVB5, a representative member of the genus *Enterovirus* in the family *Picornaviridae*, pointing to a selective inhibition of flaviviruses.

**Figure 3.**
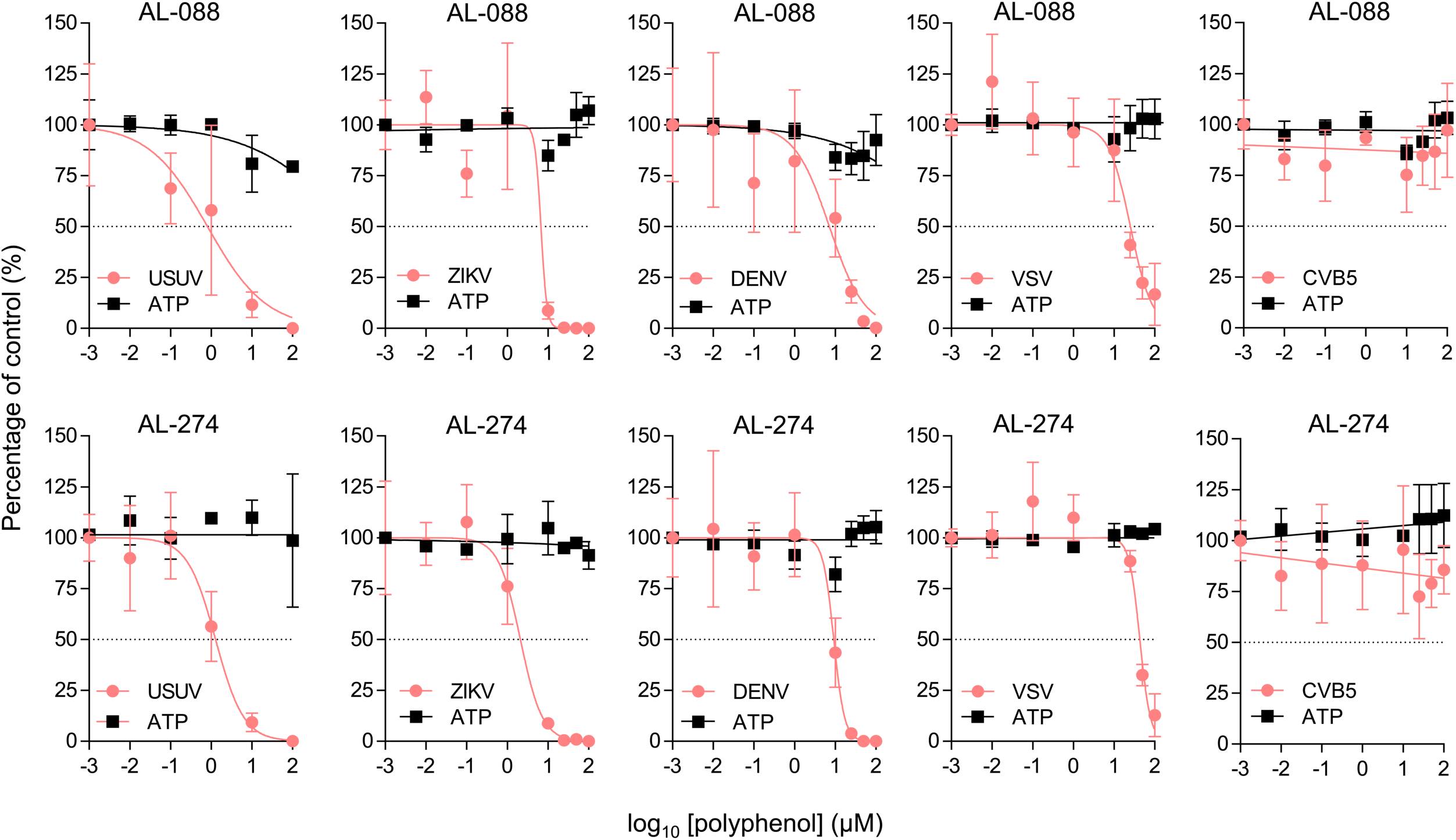
Antiviral spectrum of AL-088 and AL-274. Virus yield in the supernatant of infected Vero cells (MOI of 1 PFU/cell) was determined at 24 h pi for USUV, ZIKV, VSV and CVB5 and 48 h pi in the case of DENV-2. Polyphenols were added 1 h prior to infection and maintained throughout the rest of the assay. The cytotoxicity of the compounds was measured by determination of cellular ATP in uninfected samples at 48 h for DENV-2 and 24 h for the rest of virus. Dashed lines denote a 50% reduction. Data are expressed as mean ± SD (n = 2 to 4).

### AL-088 and AL-274 inhibit WNV multiplication and infectivity

Reporter virus particles (RVPs) produced by *trans* complementation of a subgenomic flavivirus replicon with an expression vector encoding structural proteins provide a single-cycle infectious system useful to evaluate the antiviral activity of entry inhibitors (47, 48). Accordingly, the effect of AL-088 and AL-274 on virus entry was analyzed using RVPs (42). Interestingly, pretreatment with the compounds significantly reduced the number of infected cells indicating that the compounds reduced the entry of RVPs (Fig. 4A and B). To analyze the ability of these compounds to inhibit viral multiplication, cells were infected with WNV, and treated with the polyphenols from 1 or 3 h pi (Figs. 4C and D). AL-088 and AL-274 significantly affect WNV infection when added at 1 or 3 h pi, suggesting that both compounds could also affect post-entry steps. Cells infected and treated with the polyphenols were observed by confocal microscopy (Fig. 4E). A reduction in the accumulation of both double-stranded RNA intermediates (which constitutes reliable markers of flavivirus replication) and WNV envelope (E) glycoprotein was noticed. Flavivirus replication and particle biogenesis take place coupled into the same membranous structures derived from the endoplasmic reticulum of the infected cell. Therefore, to analyze the potential effect of the treatment with the biogenesis and infectivity of the viral progeny, we compared the specific infectivity of the viral particles released form cells treated with AL-088 and AL-274 (Figs. 4F and G). The released WNV particles produced by the infected cells after treatment with AL-088 and AL-274 showed a significant reduction in their infectivity compared to vehicle (Figs. 4F and 3G), supporting that the compounds could also interfere with the morphogenesis of infectious particles. Taken together, these results suggested that galloyl derivatives exerted their antiviral effect at multiple steps of the viral replication cycle.

**Figure 4.**
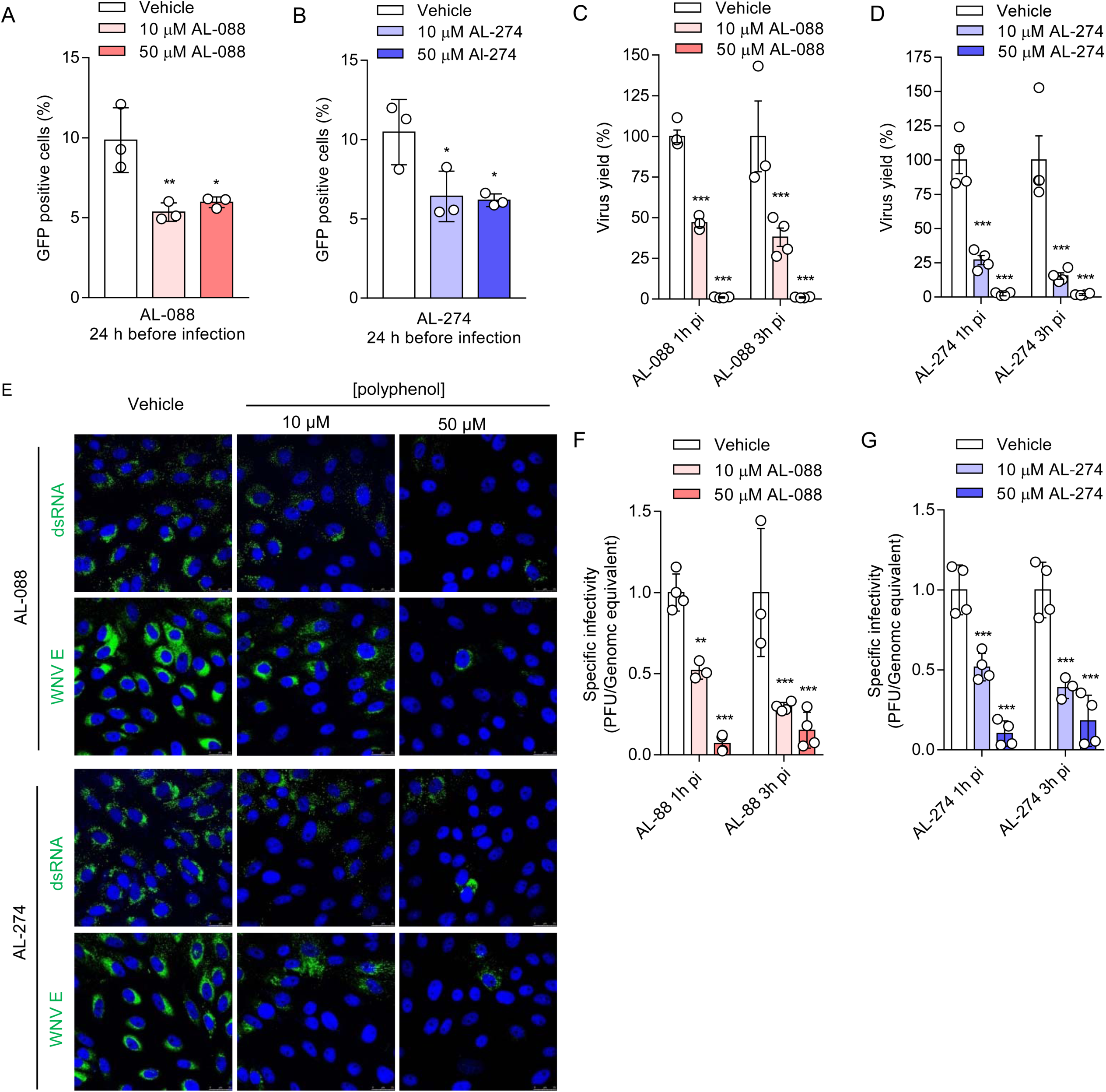
AL-088 and AL-274 inhibit WNV multiplication and infectivity. **(A and B)** Synthetic polyphenols inhibit entry of WNV RVPs. Vero cells were treated with AL-088 (**A**) or AL-274 **(B)** for 24 h and the infected with RVPs. The number of cells infected with RVPs was determined at 48 h pi by flow cytometry (n=3). **(C and D)** Synthetic polyphenols inhibit WNV multiplication when added at post-entry steps. Vero cells were infected with WNV (MOI of 1 PFU/cell) and AL-088 **(C)** or AL-274 **(D)** were added at 1 or 3 h pi. Virus yield was determined at 24 h pi (n=6). **(E)** Gallic acid derivatives inhibit the accumulation of dsRNA intermediates and viral proteins. Vero cells were infected with WNV and treated with the polyphenols from 3 h pi and fixed and processed for immunofluorescence to detect drRNA or WNV E at 24 h pi. Viral antigens are displayed in green and cellular nuclei are coloured in blue. **(F and G)** Analysis of the specific infectivity in the viral progeny released from cells treated with AL-088 **(F)** or AL-274 **(G)** as described for panels A and B (n=3 to 4). For A, B, C, D, F and G data are expressed as mean ± SD. Each dot denotes a single biological replicate. **, *P*<0.01; ***, *P*<0.001 for ANOVA and t-Student using Bonferroní’s correction.

### AL-088 and AL-274 elevate dihydrosphingomyelin levels

As already mentioned, flavivirus replication is highly dependent on the cellular lipid metabolism and previous studies supported the potential interference of AL-274 with cellular lipids (18). Therefore, we considered of particular interest to investigate whether the anti-flavivirus activity showed by AL-088 and AL-274 could be due to an alteration in the cellular lipid metabolism. With this aim, we evaluated the changes in the lipidome of cells treated with AL-088 and AL-274. In these analyses, 145 different molecular species belonging to 13 different lipid classes were identified (Fig. S1). Multivariate analyses based on the principal component analysis (PCA) indicated that both AL-088 and AL-274 altered the content of cellular lipids in a similar manner (Fig. 5A). This was evidenced by the high degree of overlay between the two groups of samples treated either with AL-088 or AL-274 that clearly differ from those treated only with vehicle (Fig. 5A). By dissecting data at the lipid subclass level, a significant increase in the amount of dihydrosphingomyelins (dhSM) was noticed in samples treated with AL-088 or AL-274 relative to control (Fig. 5B). Significantly elevated levels of dihydroceramides (dhCer) and a slight reduction of sphingomyelins (SM) were also detected in samples treated with AL-088, further supporting the effect of this polyphenol on sphingolipid metabolism. At the molecular species level, only dhSMs (d18:0/18:0, d18:0/22:0, d18:0/24:0 and d18:0/24:1) were significantly altered in samples treated with AL-088 and AL-274 (Fig. 5C). This increase can be clearly visualized in the heatmap displayed in Fig. 5D. When compared to vehicle-treated samples, the dhSM/SM ratio was significantly increased for almost all the dhSM analyzed confirming the enrichment in dhSMs in cells treated with AL-088 and AL-274 (Fig. 5E). These results indicate that these galloyl derivatives significantly alter sphingolipid metabolism and promote dhSM accumulation.

**Figure 5.**
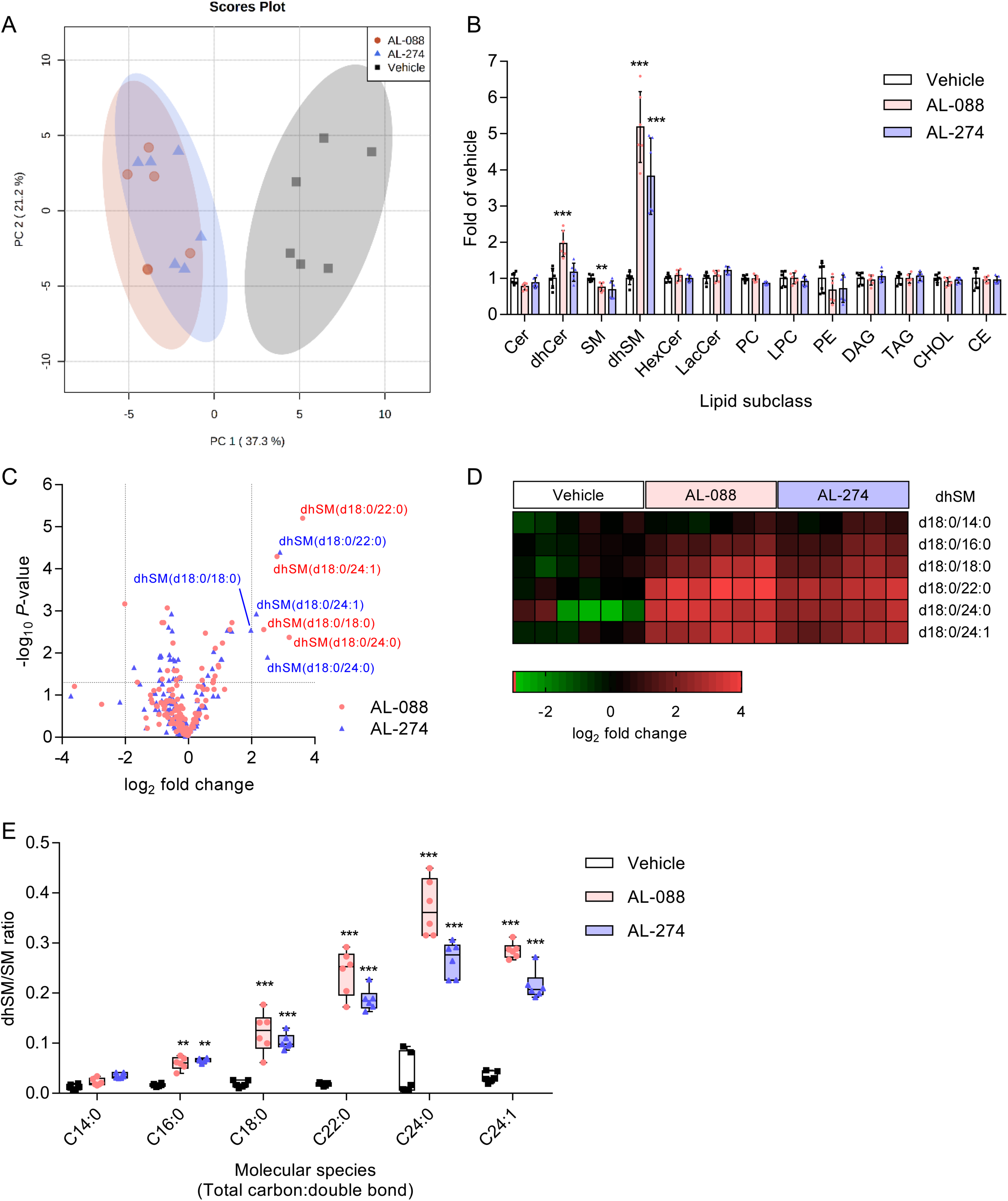
AL-088 and AL-274 elevate dihydrosphingomyelin levels. **(A)** Comparison of the lipidome of Vero cells treated with 10 μM of polyphenols (24 h) or vehicle by the PCA method. Each dot denotes a single biological replicate. 95% confidence regions for each group of samples are coloured (n=6). **(B)** Relative abundance of ceramides (Cer), dihydroceramides (dhCer), sphingomyelins (SM), dihydrosphingomyelins (dhSM), hexosylceramides (HexCer), lactosylceramides (LacCer), phosphatidylcholines (PC), lysophosphatidylcholines (LPC), phosphatidylethanolamines (PE), diacylglycerols (DAG), triacylglycerols (TAG), cholesterol (CHOL) and cholesteryl esters (CE) in samples treated with AL-088, AL-274 or vehicle. Each symbol denotes a single biological replicate (n=6). **, *P*<0.01 and ***, *P*<0.001 for multiple t-test applying Sidak-Bonferroni’s correction. **(C)** Volcano plot displaying the lipid species significantly altered in cells treated with AL-088 and AL-274. Discontinuous lines indicate a FDR *P*-value of 0.5 and log_2_fold change = 2. Each point corresponds to the mean value obtained for a single lipid species (n=6). **(D)** Heatmap displaying the relative abundance of dhSM species in samples treated with vehicle, AL-088 or AL-274. Each column denotes a single biological replicate (n=6). **(E)** Box-and-whiskers plots showing the ratio between dhSM/SM for each molecular species analyzed. Each symbol denotes a single biological replicate (n=6). **, *P*<0.01 and ***, *P*<0.001 for multiple t-test applying Sidak-Bonferroni’s correction.

### AL-088 and AL-274 inhibit ceramide desaturase

Within the sphigolipid biosynthetic pathway (Fig. 6A), the conversion of dhCer into Cer is catalyzed by sphingolipid delta(4)-desaturase (Des1) which introduces a double bond at C4-C5 in the sphingosine chain. The next step of sphingolipid synthesis is the conversion of Cer into sphingomyelin (SM) by SM synthases (SMS). In addition, SMS can also naturally convert dhCers into dhSMs, these being minor components of cellular SM (Fig. 6A). Accordingly, we hypothesized that the accumulation of dhSM in cells treated with AL-088 and AL-274 could be due to the blockage of Des1 activity. To test this hypothesis, the effect of both compounds on Des1 activity was evaluated by measuring the conversion of the fluorescent probe dhCerC6NBD into CerC6NBD, which was monitored by HPLC coupled to a fluorescence detector (45). Treatment of T98G cell cultures with AL-088 or AL-274 resulted in a reduction of dhCerC6NBD conversion, confirming that these compounds inhibited Des1 (Fig. 6B). Our compounds also inhibited Des1 in cell lysates in a dose dependent manner with IC_50_ values of 1.0 and 7.8 μM for AL-088 and AL-274, respectively (Fig. 6C). Overall, these results demonstrate that the synthetic polyphenols AL-088 and AL-274 are real inhibitors of Des1.

**Figure 6.**
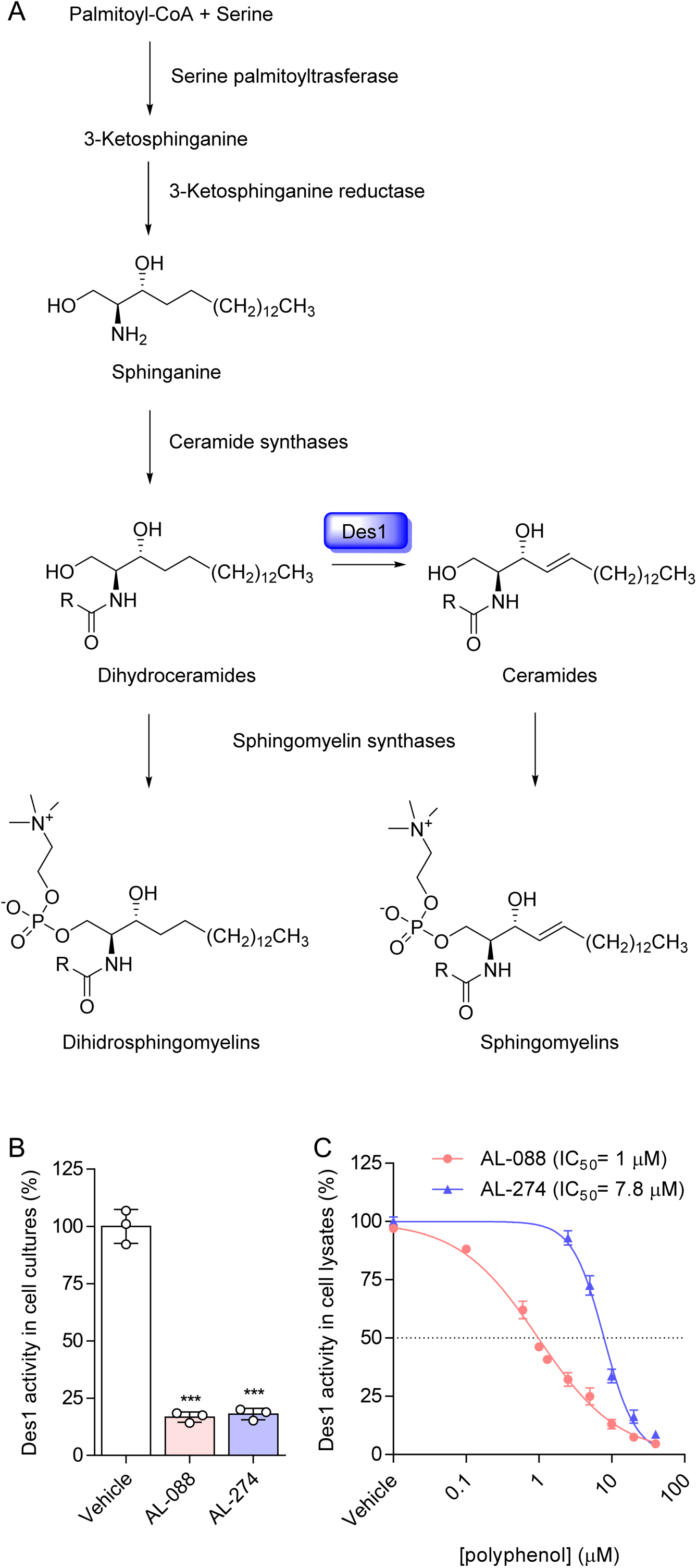
AL-088 and AL-274 inhibit ceramide desaturase activity. **(A)** Scheme of dihydrosphingolipid synthesis. Ceramide desaturase (Des1) is highlighted in blue. **(B)** Effect of AL-088 and AL-288 on the conversion of dhCerNBD to CerNBD in intact T98G cell cultures. Data are expressed as mean ± SD. Each symbol denotes a single biological replicate (n=3). ***, *P*<0.001 for ANOVA and t-Student using Bonferroni’s correction. **(C)** Dose-dependent inhibition of Des1 activity in T98G cell lysates by AL-088 and AL-274. Des1 activity was measured as described for panel B. Data are expressed as mean ± SD (n=3).

### Dihidrosphingomyelin inhibits WNV infection

The analysis of the lipidomic data pointed to the increases in dhSMs as the putative mechanism behind the antiviral activity of AL-088 and AL-274. Thus, the direct effect of this lipid on WNV infection was evaluated (Fig. 7A). The addition of exogenous dhSM significantly reduced WNV infection in a dose-dependent manner without reducing cell viability estimated by ATP measurement. This result suggests that the antiviral effect of AL-088 and AL-274 can be related to Des1 inhibition leading to the accumulation of dhSM. To further explore the potential of Des1 as a novel druggable target against WNV, the effect of the structurally unrelated Des1 inhibitor GT-11 (36) was studied (Fig. 7B). Interestingly, GT-11 inhibited WNV infection in a dose-dependent manner confirming that the pharmacological inhibition of Des1 reduced WNV infection. Overall, these results demonstrate that an increase in dhSM levels, either by adding it exogenously or by inhibiting Des1, hampers WNV replication. Moreover, these results support that Des1 is a suitable druggable target for antiviral intervention against WNV.

**Figure 7.**
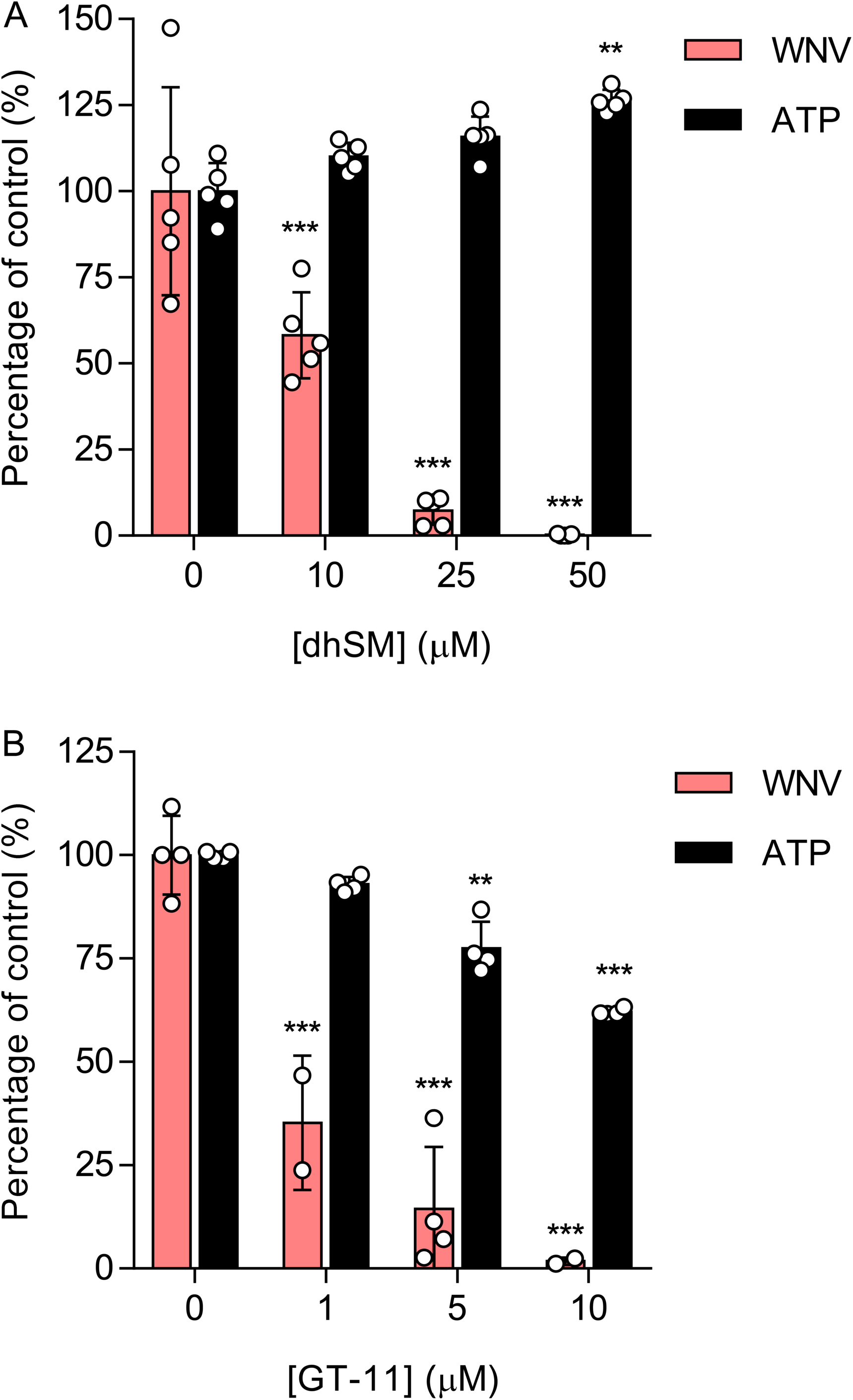
Dihydrosphingomyelin inhibits WNV infection. **(A)** Antiviral activity of dhSM against WNV. Vero cells were loaded with dhSM for 24 h, infected with WNV (MOI of 1 PFU/cell) and virus yield in the supernatant was determined at 24 h pi. The cytotoxicity of the compounds was measured by determination of cellular ATP in uninfected samples. Data are expressed as mean ± SD (n= 5). Each dot denotes a single biological replicate. **, *P*<0.01; ***, *P*<0.001 for ANOVA and t-Student using Bonferroni’s correction. **(B)** Antiviral activity of Des1 inhibitor GT-11 against WNV. Vero cells were treated with GT-11 and infected as in A. Data are expressed as mean ± SD (n = 4 - 5). Each dot denotes a single biological replicate. **, *P*<0.01; ***, *P*<0.001 for ANOVA and t-Student using Bonferroní’s correction.

## Discussion

It is well established that the amphipathic character of polyphenols, with hydrophobic aromatic rings and hydrophilic hydroxyl groups, plays a major role in their biological activities (49, 50). Incorporation of polymethylene chains may further affect this amphipathic character that can be additionally modified by functionalization of the distal part of the spacer with a second phenolic unit (18). Indeed, this approach has led in our hands to compounds with significant antiviral activity against HCV, probably affecting lipid metabolism (18). This has been now studied in detail using WNV as a clinically relevant example of flavivirus. Based on the structure-activity relation data included in the Results section, compounds AL-088 and AL-274 afforded the best selectivity indexes (>45 and >416, respectively) with EC_50_ values in the low or sub μM range against WNV (EC_50_= 2.2 and 0.24 μM, respectively in Vero cells; EC_50_=2.2 and 1.9 μM, respectively in SH-SY5Y cells). Moreover, both compounds also effectively inhibited the multiplication of other flaviviruses, namely USUV, ZIKV and DENV-2, in all cases with EC_50_ values below 10 μM. Interestingly, these compounds showed a high degree of specificity against flaviviruses when compared to other RNA viruses, exhibiting lower antiviral activity against the rhabdovirus VSV while no antiviral activity was observed against the picornavirus CVB5.

The analysis of the lipidome of Vero cells treated with AL-088 and AL-274 revealed significant changes compared to untreated cells. Particularly, a significant increase in the amount of dhSMs (d18:0/18:0, d18:0/22:0, d18:0/24:0 and d18:0/24:1) was detected, which led us to hypothesize that accumulation of dhSM could account for the antiviral effect observed with both compounds. Supporting our hypothesis, addition of external dhSM also reduced WNV infection. Moreover, when both compounds were tested as Des1 inhibitors in cells or in cell lysates, dose-dependant inhibition was detected. Indeed, similar antiviral effects were observed by treatment with the validated Des1 inhibitor GT-11. Dihydrosphigolipids, that were largely underscored since no direct biological activities could be ascribed to them and whose tissue concentrations are significantly lower than those of sphingolipids, are getting increased consideration due to their important roles in metabolic pathways and cell signalling networks (51). Their physiological roles involve, among others, response to cellular stress and autophagy, pro-death and pro-survival pathways, hypoxia or immune responses (52). However, to our knowledge, the role of dhSM in viral infections was only described for retroviruses some years ago (53–55). Key enzymes in dhSM metabolism are dhCer desaturases, with two isoforms: Des1, that is ubiquitously distributed, and Des2 that is expressed in the intestine, skin and kidney (56). A number of compounds have been described as Des1 inhibitors, that include sphingolipid analogues, such as GT-11, but also a quite variety of chemical structures, such as fenretinide (also designated as 4-HPR), Δ9-tetrahydrocannabinol (THC), the sphinosine kinase 1-2 inhibitor (SKI II) or resveratrol (56). Even more, unrevealing the mechanism of action of certain anticancer drugs, such as ABTL0812, has shown that, at least in part, their mechanism of action to induce autophagy is related to its capacity to inhibit Des1 leading to the accumulation of dhCer (57). While for GT-11, the mechanism of action involves competition with the natural substrate (36), for many other compounds, the exact mechanism of action against Des1 has not been unequivocally elucidated. As an example, fenretinide or SKI-II, which contain a 4-aminophenol in their structure have been proposed to generate reactive iminoquinones that might irreversibly react with nucleophilic residues of the protein (58). Moreover, the Fe2O3 in the active site of Des1 may contribute to oxidation of the inhibitors (58). Thus, the ability of galloyl derivatives AL-088 and AL-274 to inhibit Des1 activity, both in intact cells and cell lysates, suggests that the inhibition can occur at the protein level and/or by interacting with the Des1 associated electron transport chain, as reported for other phenolic compounds such as fenretinide, resveratrol or Δ^9^-tetrahidrocannabinol (56). Although AL-088 and AL-274 exhibited low cytotoxic profiles in Vero cells, these compounds were more cytotoxic in SH-SY5Y neuroblastoma cultures, which is consistent with previous reports on growth arrest and cytotoxicity upon Des1 inhibition in cancer cells (57, 59).

The data presented here evidence that treatment with AL-088 or AL-274 leads to an accumulation of dhSM in the host cell, which results in protection against the infection of WNV. Therefore, the accumulation of dhSM in cells treated with AL-088 and AL-274 may lead to changes in the membrane properties that result in impairment of viral infection, including inhibition of viral entry as described for human immunodeficiency virus-1 (55), or the assembly of the replication complexes and particle biogenesis. Accumulation of dhSMs has been proposed to involve the rigidification of membranes by effectively forming ordered domains through its high potential to induce intermolecular hydrogen bond (60, 61). This mechanism of action would be consistent with studies indicating that flaviviruses are highly dependent on sphigolipid metabolism and that perturbation of these pathways affects infection by targeting replication and morphogenesis (32–34, 62). In fact, the replication and biogenesis of flaviviruses take place coupled to highly remodeled cytoplasmic membranous structures derived from the endoplasmic reticulum (ER) that have a specific lipid composition (34). As these polyphenols also exerted antiviral activity against an HCV replicon system, these would further support its ability to interfere at replication stages of *Flaviviridae* (18). Thus, it is reasonable to propose that accumulation of dhSM may result in stiffening of the cellular membranes and that this severely affects multiple steps of WNV infection. The antiviral activity of AL-088 and AL-274 exerted against VSV is also consistent with previous reports indicating that sphingolipids may account for multiple roles during VSV infection. (63–65).

Our work illustrates the importance of Des1 inhibitors as host-directed antiviral agents and reveal the crucial role of dhSMs in flavivirus infection and contributes to expanding the portfolio of roles and functions of dihydrosphingolipids beyond their implication in cancer and metabolic diseases. The ability of these polyphenols to inhibit the infection of medically relevant flaviviruses makes of these compounds promising leads for the future development of antiviral therapies.

## Material and Methods

### Chemical compounds

The synthesis of the polyphenols used in this study has been previously described by us (18). *N*-[(1 *R*,2*S*)-2-hydroxy-1-hydroxymethyl-2-(2-tridecyl-1-cyclopropenyl)ethyl]octanamide (GT-11) was synthesized as described (36). Stock solutions of polyphenols and GT-11 were prepared at 10 mM in dimethyl sulphoxide (DMSO). *N*-lauroyl-D-*erythro*-sphinganylphosphorylcholine, a synthetic dhSM, was purchased form Avanti Polar lipids Inc. (Birmingham, AL) and dissolved in ethanol to perform a 50 mM stock solution. *N*-[6-[(7-nitro-2-1,3-benzoxadiazol-4-yl)amino]hexanoyl]-D-*erythro*-dihydrosphingosine (dhCerC6NBD) was synthesized as described (37).

### Viruses and infections

Virus infections were performed on Vero CCL-81 or SH-SY5Y CRL-2266 cells (ATTC, Manassas, VA). Vero cells were cultured in MEM (Corning, Manassas, VA) supplemented with 10% fetal bovine serum (Gibco, Life Technologies, Paisley, UK) and SH-SY5Y cells were grown in DMEM-F12 (Biowest, Nuaillé, France) supplemented with 10 % fetal bovine serum. Penicillin-streptomycin mixture (Corning) and L-glutamine (Corning) were also added to cell cultures. WNV New York 99 (38), USUV SAAR 1776 (38), American ZIKV PA259459 (39), DENV-2 (16), vesicular stomatitis virus (VSV) Indiana (39) and Coxsackievirus B5 (CVB5) strain Faulkner (40) were used. Procedures for infections in liquid medium and virus titrations in Vero cells in semisolid agar medium have been previously described (40, 41). WNV, USUV, ZIKV, VSV and CVB5 titers were determined 24 h postinfection (pi). DENV-2 titers were determined 48 h pi. A multiplicity of infection (MOI) of 1 PFU/cell was used for all experiments, except when indicated.

### Cellular toxicity

Cell viability was measured in uninfected cells by ATP quantification using the Cell Titer Glo luminescent cell viability assay (Promega, Madison, WI). Proliferation was measured after 24 h of treatment of uninfected cells plated at low density (<50% of confluence) by determination of the cell number with a TC20 automated cell counter (Bio-Rad, Hercules, CA) and Trypan Blue dye (Bio-Rad).

### Reporter virus particles (RVPs)

WNV single-round reporter virus particles (42) were produced by complementation in *trans* of a sub-genomic reporter replicon including GFP (kindly provided by T.C. Pierson) with WNV structural proteins using a pcDNA 3.1 (+) expression vector that encoded the prM and E proteins of WNV Novi Sad/12 (GenBank: KC407673.1). Plasmids were transfected into human embrionic kidney (HEK) 293-T cells (ATCC CRL-11268) with DharmaFECT kb DNA transfection reagent (Dharmacon, Lafayette, CO). The RVPs in the supernatant were harvested at 48 h post-transfection and used to infect Vero cells. The amount of Vero cells expressing GFP was determined by flow cytometry using a FACSCanto II (BD Biosciences, Erembodegem, Belgium) at 48 h pi.

### Specific infectivity

Specific infectivity was calculated as the ratio between PFU determined by virus titration and PFU equivalents determined by quantitative RT-PCR by comparison with previously titrated standards. RNA extraction and one-step reverse transcription coupled to quantitative PCR were performed as described (30).

### Immunofluorescence and confocal microscopy

Mouse monoclonal antibody J2 against double-stranded RNA (dsRNA) was from Scicons (Budapest, Hungary). Mouse monoclonal antibody MAB8150 against WNV E glycoprotein was from EMD Millipore (Billenca MA). Alexa Fluor 488 goat anti-mouse IgG (H+L) and TO-PRO-3 were from Invitrogen. Immunofluorescence and confocal microscopy were performed as described (40).

### Lipidomics

Lipid extractions, identification, and quantification by mass spectrometry of Vero cells (10^6^ cells/determination) treated with the inhibitors (10 μM, 24 h), or drug vehicle, were performed as described (32, 43). Fold change in lipid levels between control and treated cells was calculated as log_2_ (treated/control). The principal component analysis (PCA) was performed with Metaboanalyst 5.0 (44).

### Dihydroceramide desaturase activity

Desaturase activity was measured in T98 intact cells or lysates after 4 h of incubation with the polyphenols by quantification of the conversion of the dhCerC6NBD probe into CerC6NBD substrate by HPLC coupled to a fluorescence detector (45).

### Data analysis

Data are presented as mean ± standard deviation (SD). The number of independent biological replicates (*n*) analyzed is indicated in the figure legends. Prism 7 for Windows (GraphPad Sofware, San Diego, CA) was used for statistical analyses. Dose-response curves were calculated by adjusting the sigmoidal log (inhibitor) *vs* normalized response (variable slope) or linear regression in the case of CVB5. Means were compared using one-way analysis of the variance (ANOVA) corrected for multiple comparisons with Bonferroni’s correction for pairwise comparisons of multiple treatment groups with a single control group. In the case of lipidomic analyses, comparisons were performed using multiple Student’s t-test and Sidak-Bonferroni correction for lipid subclasses and the false discovery rate (FDR) for lipid species. Significantly altered lipids were considered when *P*<0.05 and log2 fold change over vehicle >1.5. Statistically significant differences are denoted by asterisks. *, *P*<0.05; **, *P*<0.01; *** *P*<0.001).

## Acknowledgements

We thank Theodore C. Pierson (National Institutes of Health, USA) for the subgenomic replicon of WNV. This work was supported by Spanish Ministry of Science and innovation AEI / 10.13039/501100011033 under grant PID2019-105117RR-C21 (to MAM-A); PID2019-105117RR-C21 (to M-JP-P); and PID2020-119195RJ-I00 (to N-JO); and from AECSIC under grant PIE-201980E100 (to M-JP-P and AS-F). This research work was also funded by the European Commission – NextGenerationEU (Regulation EU 2020/2094), through CSIC’s Global Health Platform (PTI Salud Global). PM-C was supported by a FPI fellowship (PRE2020-093374) from AEI / 10.13039/501100011033. The funders had no role in study design, data collection and interpretation, or the decision to submit the work for publication.

